# Practically Error-Free Junctions Enable Solving Large Instances of Exact Cover Problems Using Network-Based Biocomputation

**DOI:** 10.1101/2025.09.10.675367

**Authors:** Eugene Christo V R, Christoph Robert Meinecke, Bert Nitzsche, Roman Lyttleton, Cordula Reuther, Danny Reuter, Heiner Linke, Till Korten, Stefan Diez

## Abstract

Network-based biocomputing (NBC) presents an energy-efficient, parallel computing approach for solving nondeterministic polynomial time (NP) complete problems by leveraging motor-driven cytoskeletal filaments that explore all possible solutions through nanofabricated networks in a massively parallel fashion. However, guiding errors at pass junctions, where filaments deviate from their intended path, currently limit the scalability of NBC systems. In this study, we addressed this critical challenge by fabricating sub-200 nm channel geometries using modified electron-beam-lithography and reactive-ion-etching protocols to physically constrain the trajectories of kinesin-driven microtubules and enhance path fidelity. Investigating junction designs with varying channel widths, we demonstrate that reducing channel width significantly lowers junction error rates. Practically error-free junction performance was achieved by scaling down the entire network geometry by a factor of two. These optimized junctions were incorporated into NBC networks that successfully solved 24- and 25-set instances of the Exact Cover problem, representing solution spaces of approximately 16 million and 33 million, respectively. This work establishes a new benchmark in NBC performance and represents a computational scale far beyond what has been achieved in prior demonstrations.

## Introduction

The increasing prevalence of data-intensive applications in areas such as operations research, molecular biology, cryptography, and artificial intelligence has emphasized the need for solving complex combinatorial problems ^1–5^. A particularly important class of such problems is those categorized as non-deterministic polynomial-time complete (NP-complete) problems. Their computational complexity grows exponentially with input size, rendering them intractable on conventional, serial operating computers. Although heuristic and approximate algorithms can sometimes reduce computation time, they often sacrifice accuracy and cannot guarantee a solution within acceptable time frames for all instances ^6,7^. Furthermore, the stagnation of Moore’s Law and the rising energy losses associated with continued CMOS miniaturization have spurred interest in exploring alternate, energy-efficient computing approaches ^8–10^.

In this context, network-based biocomputation (NBC) has emerged as a promising parallel computing strategy that leverages the nanoscale transport properties of molecular motors and cytoskeletal filaments in engineered synthetic environments ^11^. Here, a combinatorial problem is physically encoded into a nanofabricated network, and kinesin-driven microtubule filaments solve the problem by exploring all potential paths simultaneously. NBC devices require orders of magnitude less energy than traditional electronics due to the inherent energy-efficiency of molecular motors and the fundamental energy advantage of parallel computing ^12^.

In this study, we use the Exact Cover problem, relevant to resource-allocation tasks such as airline fleet planning ^2^ and cloud-computing scheduling ^5^, both as a benchmark problem and as an example for illustrating the operation of NBC. The Exact Cover problem consists of a target set X and a collection of subsets S. The goal is to determine whether there exists a subcollection S^*^⊆ S such that every element of X appears exactly once across the subsets in S^*^. In other words, the chosen subsets S^*^ must be pairwise disjoint, and collectively cover all the elements of target set X without omissions or repetitions.

The implementation of Exact Cover on an NBC device (**Figure 1 A-C**), as described in our previous works ^13^, involves mapping each element of the set *X* to a binary position. Each subset in *S* is similarly represented as a binary number according to this mapping, where a bit set to ‘1’ indicates the presence of the corresponding element in *X*, and a ‘0’ its absence. The numerical values assigned to the elements of *X* are arbitrary and hold no computational significance; only their mapping positions matter. For instance, whether the elements are numbered 1–6 or randomly between 115–12849, the computation proceeds identically, since each element is uniquely mapped to a bit position. The number of required binary position required is solely determined by the size of the target set *X*.

**Figure 1.**
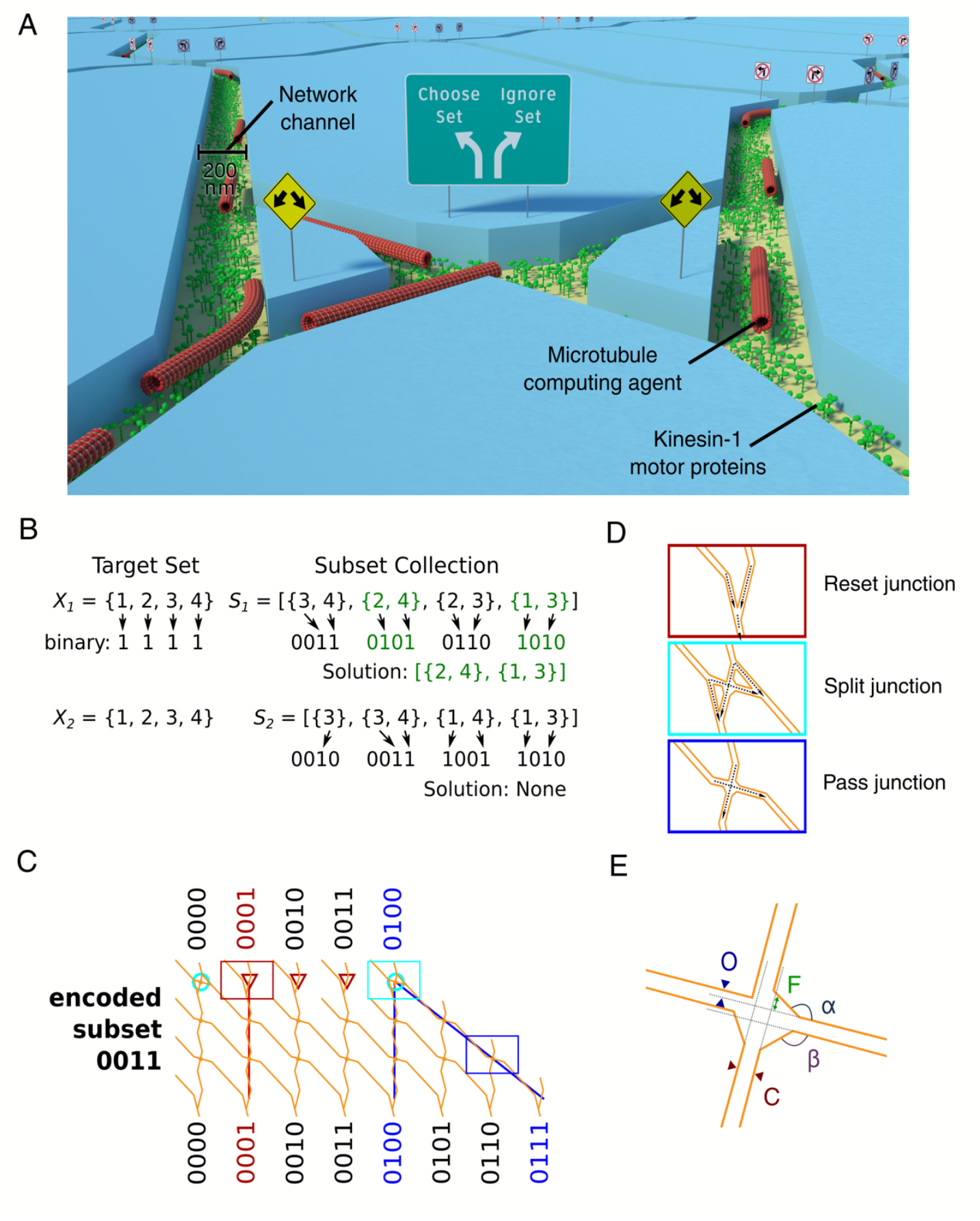
Working principle of an NBC network that encodes Exact Cover. (**A**) Schematic rendering of a biocomputation network. Kinesin-1 motors coat the bottom of 200 nm wide channels etched in poly (methyl methacrylate) (PMMA). The channels are explored by microtubules propelled by kinesin-1 motors. (**B**) Binary encoding principle: Elements of the target set X are mapped to bits of a binary number. Each subset in S is converted to a binary number by the same mapping rule. i.e., a bit is set to 1 if the respective element in X is a member of the subset and set to 0 otherwise. Two examples of exact cover instances (Xa Sa and Xb Sb) are given. Xa Sa has a solution (highlighted in green) and Xb Sb does not. (**C**) Example network block that encodes the subset 0011. Microtubule agents arriving at input 0100 (blue) encounter a split junction (example denoted by cyan rectangle) that allows the agents to randomly choose (diagonal path) or disregard (the path straight down) the subset encoded by this network block. If the subset is chosen, the number 0011 is added to the input 0100 and the agent arrives at the output 0111. Otherwise, the agent stays in the input column 0100. Because input subsets containing identical elements cannot be combined (as this would violate the rules of exact cover), all inputs that contain elements already present in the encoded subset start with a reset junction (example denoted by red rectangle). This junction forces the agents to take the path straight down, i.e. disregarding the encoded subset. For example, an agent entering at input 0001 (red rectangle) can only leave at output 0001. The input and output rows are separated by two rows of pass junctions (example denoted by blue rectangle) that allow agents to continue a diagonal path, incrementing the value of the binary number, or continue moving directly down and maintaining the value of the binary number. Thus, the number of pass junction rows ultimately defines which elements are contained in the encoded subset. A full network can contain many network blocks. (**D**) Schematic representation of each junction type. The color of the frame corresponds to the rectangles in (C). The black dotted arrows represent the permissible microtubule paths. (A-D) Adapted with permission under a Creative Commons Attribution 4.0 license from Korten et. al ^13^. Copyright 2021 IOP Publishing. (**E**) Schematic of pass junction illustrating its geometric parameters such as channel width (denoted as C), offset (denoted as O), funnel width (denoted as F), and funnel angles (denoted as α and β). Adapted from Surendiran, P et. al ^16^.

Each binary subset defines a network block that guides microtubule trajectories through three distinct junction types. Microtubule agents arriving at a split junction are allowed to randomly choose (diagonal path) or disregard (the path straight down) the subset encoded by this network block. While microtubule agents encountering a reset junction are forced to take the path straight down, i.e disregarding the encoded subset. Because input subsets containing identical elements cannot be combined (as this would violate the rules of exact cover), all inputs that contain elements already present in the encoded subset encounter a reset junction. The input and output rows are separated by pass junctions that allow agents to continue a diagonal path, incrementing the value of the binary number, or continue moving directly down and maintaining the value of the binary number. The permitted path a microtubule can take in these junctions are denoted by black dotted arrows in **Figure 1D**. All the network blocks are concatenated to form the full computational network. The goal is for a microtubule agent to reach a final binary state where all bits are set to ‘1’, corresponding to a complete non-redundant cover for target set *X*. The presence of such a path indicates a valid Exact Cover solution. All paths are explored simultaneously and the computational power of NBC lies in this massively parallel exploration capability. Proof-of-concept studies have demonstrated NBC’s ability to solve small instances of NP-complete problems, including the Subset-Sum Problem (SSP) with eight possible solutions ^11^, the Exact Cover problem with 1024 possible solutions ^14^, and 3-Satisfiability (3-SAT) problems involving up to three variables and five clauses ^15^.

However, further scaling NBC to solve larger and more practically relevant combinatorial problems is currently hindered by the error rates associated with NBC. One principal source of error is junction errors, which occur at pass junctions when filaments deviate from their intended paths and take a wrong turn, leading to incorrect computations. As junction errors scale with network size ^16,17^, they become increasingly significant for larger combinatorial problems, since the number of pass junctions a microtubule must traverse increases proportionally. Consequently, the probability that a microtubule follows the correct path across all junctions scales as (1 – *E*)^*x*^, where *x* is the number of junctions crossed. For instance, with a junction error rate of approximately 2%, networks with only tens of rows can be reliably explored. In contrast, error rates of 0.5% and 0.1% allow the use of networks with hundreds and thousands of rows, respectively. This highlights the critical impact of even small junction error rates on the overall computational fidelity of the NBC systems. Therefore, minimizing junction errors, which is the focus of this work, is essential in order to solve large combinatorial problems of practical importance ^16^.

Here we demonstrate a substantial increase in the computational scale of NBC, enabled by practically error-free operation of pass junctions. We used a stepwise approach, where we first designed a series of junction geometries with varying channel widths, deliberately using an error-prone junction design to quantify how geometric confinement influences computational fidelity. Consistent with prior experimental and simulational studies ^16,18,19^, we found that reducing the channel width constrains filament trajectories, promotes straighter microtubule gliding, and consequently lowers junction error rates. The devices were produced using electron-beam lithography combined with reactive-ion etching to create sub-200 nm channels compatible with kinesin binding and microtubule motility. Building on this, we achieved practically error-free operation of pass junctions using optimised geometry from prior work, along with further downscaling of the entire network. Finally, we incorporated the optimized junction design into functional NBC devices encoding 24-set and 25-set instances of the Exact Cover problem. These devices successfully solved problem instances comprising approximately 16 million and 33 million potential solutions, respectively, establishing a new benchmark for NBC performance.

## Results

### Design and Fabrication of Junction optimization devices

We first investigated the effect of channel width on guiding filament trajectories within the pass junction. Although channel width is a key parameter, the overall geometry of the pass junction is more complex. Additional parameters such as the offset distance, funnel width, and funnel angles α and β (**Figure 1E**), also significantly influence junction performance ^16^.

For experimental simplicity, we selected a junction design known to exhibit an inherently higher error rate ^16^. These junctions had an offset distance of 0 nm, a funnel width of 66 nm, and funnel angles α = 45° and β = 30°, which were kept constant, while the channel width was varied from 50 nm to 200 nm. As a first step, these junction designs were incorporated into four small computational networks (**Figure 2A**) for testing purposes. Each network consisted of a landing zone where microtubules could attach and subsequently be transported into the computation area, where they explored the networks comprised of pass and split junctions. To extend the operational lifetime of filaments within the devices, feedback loops and rectifiers enabled continuous recycling of microtubules and ensured robust unidirectional gliding of microtubule agents.

**Figure 2.**
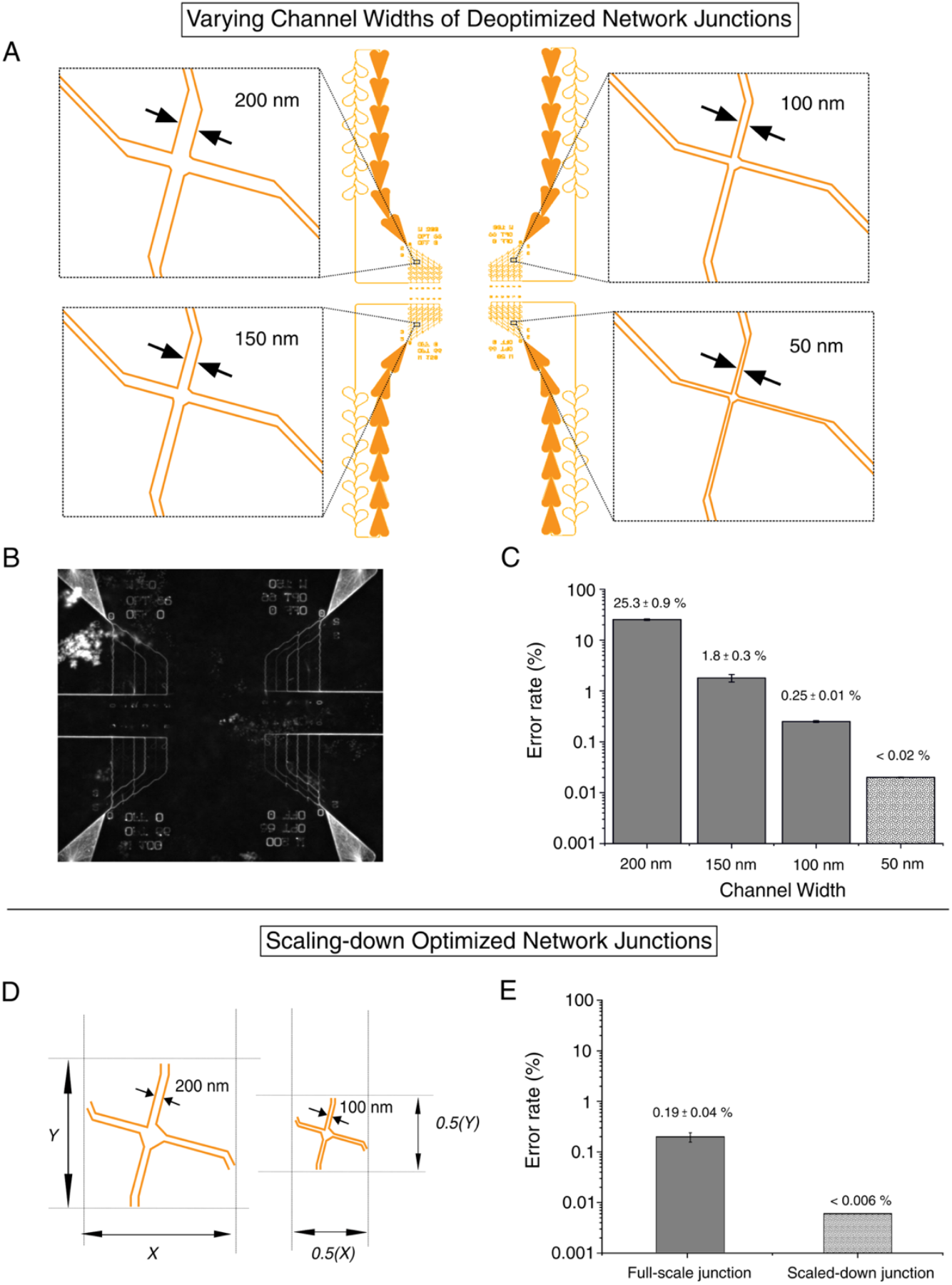
Optimization of pass junction geometry to minimize junction error rates. (**A**) Schematic of junction optimization networks layout showing the design of pass junctions with varying channel widths. The channel widths were varied between 50 nm to 200 nm. Inset: 200 nm channel is shown in top left box, 150 nm channel is shown in bottom left box, 100 nm channel is shown in top right box and 50 nm channel is depicted in bottom right box. (**B**) maximum projection of microtubule tracks along the fabricated networks. (**C**) Bar plots showing the error rates with respect to the channel widths. Error bars denote the counting errors (square root of the values). (**D**) Schematic illustration of full-scale and scaled-down junction designs. (**E**) Bar plots showing the error rates for the full size (200 nm channels) and scaled down (100 nm channel) junctions.

The fabricated junctions (**Methods** and **Supplementary Figure S1**) were characterized using scanning electron microscopy (SEM) and atomic force microscopy (AFM) (**Supplementary Figure S2**). To evaluate the performance of these junctions, kinesin-coated networks were then explored by fluorescently labeled microtubules (**Methods**). **Figure 2B** shows the maximum projection of microtubule paths across the networks. All microtubule paths at pass junctions were analyzed to quantify the junction error rates (E), defined as

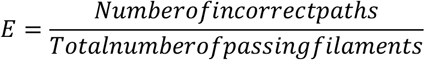

The measured junction error rates (see **Methods**) demonstrated an inverse relationship with channel width. Specifically, 200 nm wide channels exhibited an error rate of 25.3% ± 0.9% (mean ± standard deviation, n = 3,022), while 150 nm channels showed a significantly reduced error rate of 1.8% ± 0.3% (n = 2,548). Narrowing the channels to 100 nm yielded an error rate of 0.25% ± 0.01% (n = 2,444. For 50 nm-wide channels, no junction errors were observed across 5122 events, indicating an error rate below 0.02% (**Figure 2C**).

Building on these findings, the design improvements were applied to the optimized junctions currently used in state-of-the-art NBC devices ^14^ to further reduce their junction errors. For the improved design, the entire network geometry comprised of 200 nm wide channels was uniformly scaled down by a factor of two in both length and width dimensions leading to a network with 100 nm channels throughout the network (**Figure 2D**). We chose a final channel width of 100 nm, not the 50 nm that had led to the lowest error in the small scale experiments, to improve the overall performance of the device. I.e. to avoid too small loading zones, to reduce the risk of blocked channels and to improve the number of microtubules able to glide in a channel in parallel. The full-scale pass junctions exhibited an error rate of 0.19% ± 0.04% (n = 11,878), whereas the scaled-down junctions showed no errors across 17,379 events, corresponding to an error rate of less than 0.006% (**Figure 2E**). To demonstrate scale up of the technique, this practically error-free junction design was incorporated into computational networks to solve a 24-set and 25-set instance of the Exact Cover problem.

### Encoding 24-set and 25-set Exact Cover Instances on to NBC devices

Both the 24-set and 25-set instances of the Exact Cover problem used the same six-element target set, *X* = {1, 2, 3, 4, 5, 6} but differed in their subset collections and whether a valid solution existed. The 25-set instance included a collection, *S*_*1*_ = [{2,1}, {3,1}, {3,2,1}, {4,2}, {6,3,2}, {6,3,2,1}, {6,4}, {6,4,1}, {6,4,2}, {6,4,2,1}, {6,4,3}, {6,4,3,1}, {6,4,3,2}, {6,4,3,2,1}, {6,5}, {6,5,1}, {6,5,2}, {6,5,2,1}, {6,5,3}, {6,5,3,2}, {6,5,3,2,1}, {6,5,4,1}, {6,5,4,2,1}, {6,5,4,3,1}, {6,5,4,3,2}]. This instance was designed to have exactly one valid solution, consisting of the subsets [{3,1}, {4,2}, {6,5}] that together cover all elements of *X* exactly once and without repetition. In contrast, the 24-set instance involved a modified collection, identical to *S*_*1*_ but with a critical subset {6,5} omitted: *S*_*2*_ = [{2,1}, {3,1}, {3,2,1}, {4,2}, {6,3,2}, {6,3,2,1}, {6,4}, {6,4,1}, {6,4,2}, {6,4,2,1}, {6,4,3}, {6,4,3,1}, {6,4,3,2}, {6,4,3,2,1}, {6,5,1}, {6,5,2}, {6,5,2,1}, {6,5,3}, {6,5,3,2}, {6,5,3,2,1}, {6,5,4,1}, {6,5,4,2,1}, {6,5,4,3,1}, {6,5,4,3,2}]. In this instance, no subcollection satisfied the exact cover condition and therefore no valid solution existed.

To efficiently encode these large instances while minimizing network complexity, we employed a previously developed strategy known as reverse exploration ^13^. Since the target exit represents a binary string with all bits set to ‘1’, the network was split into a forward and reverse section, each encoding only half the total number of sets represented in the network. The forward network is explored from designated entrances and encodes the first portion of the subset collection, while the reverse network is explored backward from the target exit, and encodes the remaining subsets. The solution is identified when microtubules from both directions converge at an intermediate node, representing a valid subset collection that cover the target set. This approach reduces network size, lowers the number of required filaments and shortens the time needed to find a solution. For the 25-set instance, the exploration paths converge along the path corresponding to the correct solution subsets [{3,1}, {4,2}, {6,5}] (**Figure 3A**, the permitted microtubule paths indicated in blue), while for the 24-set instance, no such convergence exist, indicating the absence of a valid solution (**Figure 3B**, the permitted microtubule paths indicated in blue).

**Figure 3.**
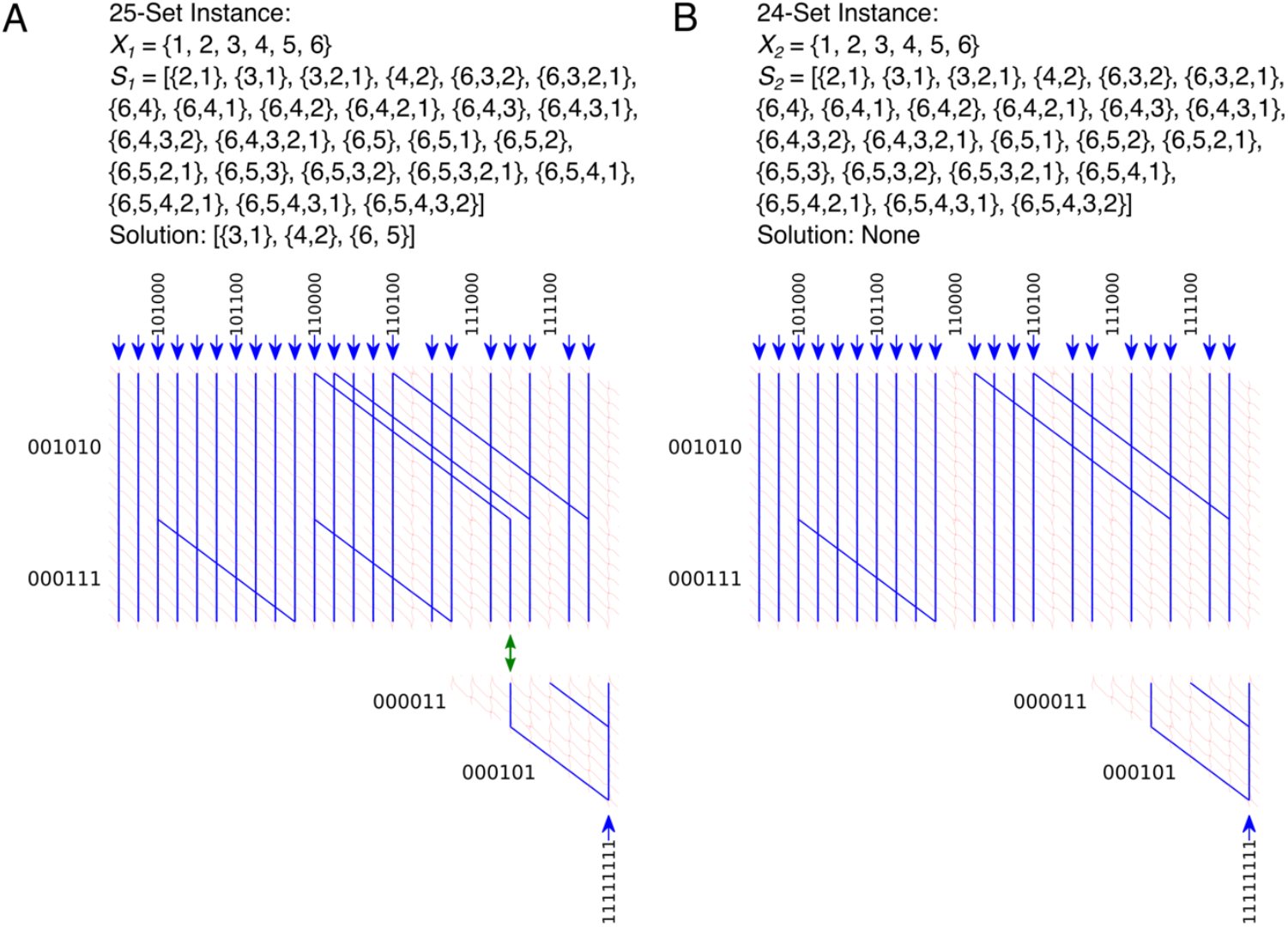
Layouts of reverse Exact Cover devices to solve Exact Cover problems with 25 and 24 set instances (33 and 16 million possible solutions). (**A**) Network layout of the Exact Cover instance with 25-sets corresponding to the binary numbers: 000011, 000101, 000111, 001010, 100110, 100111, 101000, 101001, 101010, 101011, 101100, 101101, 101110, 101111, 110000, 110001, 110010, 110011, 110100, 110110, 110111, 111001, 111011, 111101, 111110. This Exact Cover instance has a solution, which is indicated by the green double arrow. (**B**) Network layout of the Exact Cover instance with 24-sets corresponding to the binary numbers: 000011, 000101, 000111, 001010, 100110, 100111, 101000, 101001, 101010, 101011, 101100, 101101, 101110, 101111, 110001, 110010, 110011, 110100, 110110, 110111, 111001, 111011, 111101, 111110. This Exact Cover instance has no solution. (A, B) Correct paths are highlighted in blue.

### Solving 24 and 25 set instances of Exact Cover problem with NBC

The networks were explored by fluorescently labeled, kinesin-propelled microtubules. Filament movement was tracked using time-lapse fluorescent microscopy with a 2-second interval. To assess the computational output, the number of microtubule filaments reaching each network exit were statistically analyzed. The statistical significance of each outcome was determined as previously described ^14^ (see also **Methods**). The filament counts at each exit are shown in **Figure 4C-F**. Exits where filament counts exceeded a predefined significance threshold (p < 0.05, green dash-dotted lines) were classified as correct exits (shown in green) Exits below a corresponding lower threshold (magenta dotted lines) were classified as incorrect (shown in red).

**Figure 4.**
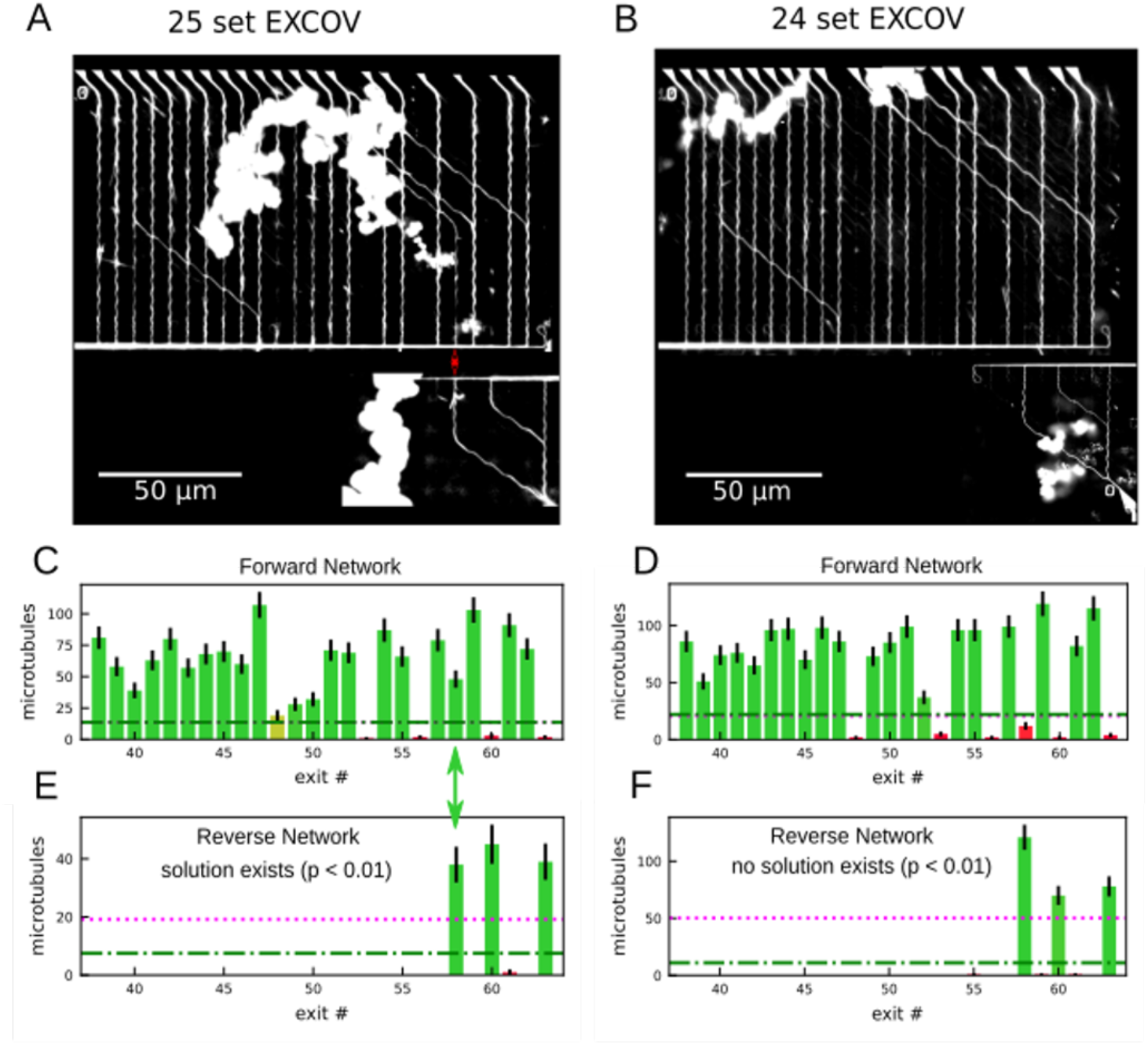
Microtubule results for the 25-set and 24-set Exact Cover instances: (**A, C, E**) 25-set instance with exactly one solution (indicated by green double arrow in C and E). (**B, D, F**) 24-set instance without solution. (A, B) Standard deviation plots of 600 frames of 2 s time-lapse fluorescence micrographs of microtubules moving through the computational network. Paths that were more frequently visited by microtubules are brighter. (C, D, E, F) Number of microtubules counted at each exit of the (C, D) forward- and (E, F) reverse halves of the Exact Cover networks. RGB colors indicate probabilities that the respective counts correspond to a correct (more green) or an incorrect (more red) exit. Error bars represent the counting error (square root of the respective value). Values above the green dash-dotted lines are significant correct exits (p < 0.05), values below the magenta dotted lines are significant incorrect exits (p < 0.05). In total, the calculation took 20 min for each network.

During NBC operation, no pass junction errors were observed across 17,379 events, confirming the reliability of the optimized junction design. This practically error-free performance enabled microtubules to traverse the networks with an overall path success probability exceeding 99.9%. Given that each microtubule traverses 17 pass junctions, even a single error (error rate ∼ 0.000057) would reduce the overall path success probability to approximately 99.9 % ((1−0.000057)^17^). Since, no such errors were observed, the actual path success probability is even higher.

Despite this high fidelity of pass junctions, a few microtubule filaments exited the network incorrectly due to landing errors, where microtubules landed at unintended positions within the network rather than through the designated entrances. Out of 2667 total exit events, 127 events were attributed to such errors. These included some filaments that still exited at correct outputs and others that exited at unintended ones. Nevertheless, the most frequently traveled microtubule paths closely matched the prediction by the network design for valid solutions (**Figure 4A and B**).

For the 25-set instance, microtubules from the forward and reverse networks successfully converged at the correct solution path at exit point 111010, confirming the existence and successful identification of a valid Exact Cover (p<0.01; significance was determined as described in ^14^). This solution followed a complete path starting at subset {5,6}, corresponding to entrance 110000, proceeding through subset {2,4} (001010) in the forward network and through subset {1,3} (000101) in the reverse network, forming a path to the target set {1,2,3,4,5,6} (111111). In contrast, the 24-subset instance showed no such convergence between forward and reverse paths, correctly indicating that no solution exists for that Exact Cover instance (p<0.01).

While narrower channels effectively reduced junction errors, they raised the question of whether reduced kinesin density in narrower channels might increase microtubule detachment at network junctions. To assess this, detachment rates were experimentally compared between networks fabricated with 100 nm and 200 nm-wide channels (full-scaled and scaled down 25-set Exact Cover networks). No significant difference in the detachment rate per junction was observed between the two networks. In the full-scale device, 10 out of 42 microtubules filaments detached before crossing 17 pass junctions, while 11 out of 55 microtubule filaments detached before crossing 17 pass junctions in the scaled down device (**Supplementary Figure S3**).

The percentage of microtubules traversing each junction as a function of the number of junctions traversed was fitted using linear regression, assuming a first-order approximation of the exponential decay function, for both networks. The slope for the full-scale network was –1.27 ± 0.11 per junction, and for the scaled-down network was –1.41 ± 0.11 per junction. Both networks showed significant negative correlations between junction number and detachment rate. Statistical comparison using an interaction model revealed no significant difference between slopes (p = 0.372), indicating similar detachment kinetics across network scales (**Figure 5**).

**Figure 5.**
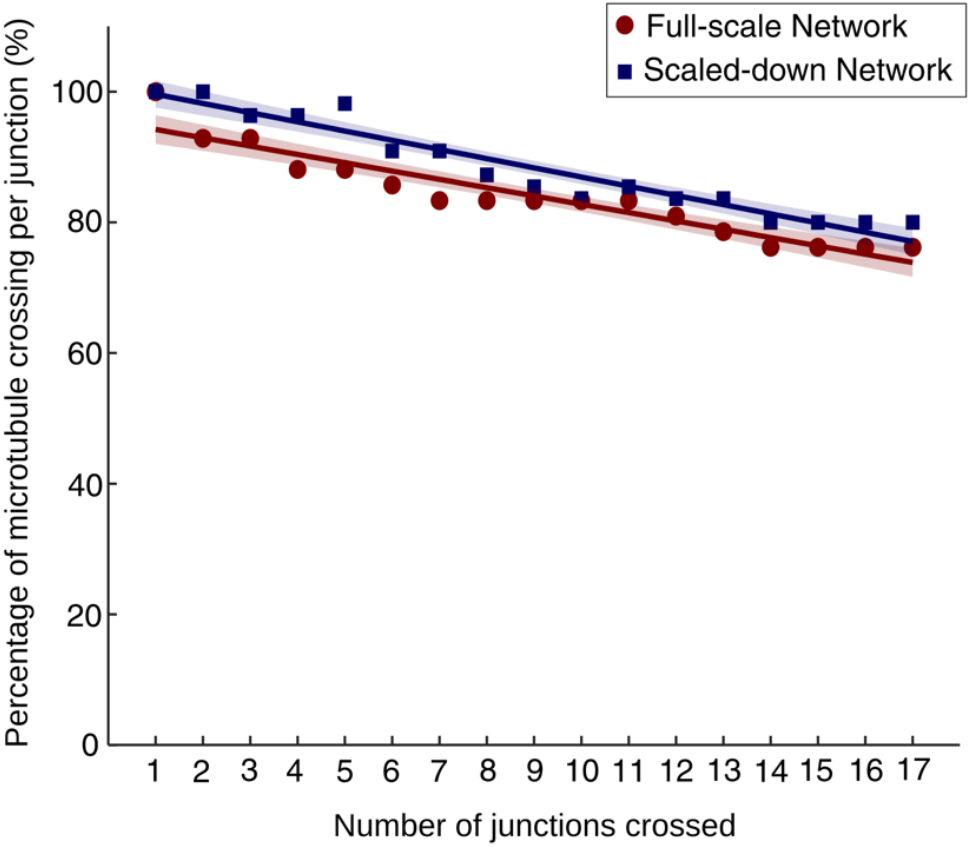
Detachment rate of microtubules per junction. Percentage of microtubules traversing each junction as a function of the number of junctions crossed in the full-scale network (red circles) and scaled-down network (blue squares). Lines represent linear regression fits for each condition, with shaded areas indicating 95% confidence intervals. The slope for the full-scale network was –1.27 ± 0.11 per junction, and for the scaled-down network was –1.41 ± 0.11 per junction. Both networks show significant negative correlations between junction number and detachment rate. Statistical comparison using an interaction model revealed no significant difference between slopes (p = 0.372), indicating similar detachment kinetics in both networks.

These experiments demonstrate the successful design and implementation of NBC devices capable of solving larger instances of the Exact Cover problem, specifically, a 25-set instance (with a known solution and a sample space of approximately 33 million) and a 24-set instance (with no solution and a sample space of approximately 16 million). The integration of reverse exploration, binary encoding, and practically error-free junctions enabled NBC systems to explore these vast combinatorial spaces in parallel with high efficiency and accuracy. Solving these problem instances marks a significant advancement in molecular computing, representing a new benchmark for NBC based computing.

## Discussion

In this study, we addressed junction errors, a key limitation hindering the scalability of NBC systems. These errors become increasingly significant as network complexity increases, since the number of junctions a microtubule must traverse and thus the probability of junction error, grows with problem size. To minimize these errors, we aimed to physically constrain filament trajectories and improve path fidelity. Toward this end, a modified electron beam lithography and optimized reactive ion etching protocol was used to fabricate sub-200 nm channel geometries compatible with microtubule gliding assays. We then used junction designs known to exhibit high error rates to demonstrate that reducing channel width significantly lowers junction errors. Narrower channels increase the angular deviation required for a gliding microtubule to make an unintended turn, effectively preventing such errors (**Figure 6**).

**Figure 6.**
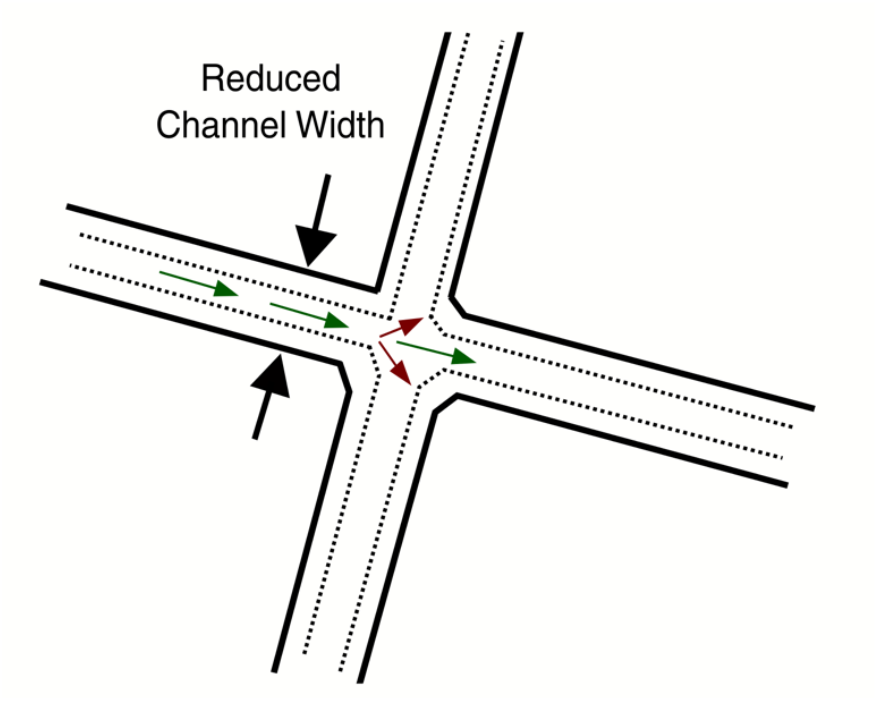
Schematic of microtubule path in a pass junction. Movement of the microtubule in the correct path is represented by green arrows. The red arrows represent a wrong turn. The black arrows show the reduced width of the channel, represented by the dotted lines. Microtubules have to turn along steeper angles to make a wrong turn in narrower channels.

The improved fabrication technology enabled us to scale down the state-of-the-art junction design, which exhibited an error rate of 0.19%. In the scaled-down networks, we did not observe a single error across 17,379 junction-crossing events. Unlike three-dimensional junctions with overpasses ^17^, the design used here does not eliminate errors in principle; however, in practice, all pass junctions operated error-free. Importantly, this design is considerably simpler to fabricate, relying on established lithography methods. The reduction in fabrication steps not only streamlined production but also improved junction smoothness by removing residual PMMA monolayers from the channel floor, thereby enhancing performance. In addition to improving fidelity, down-scaling reduced microtubule exploration distances, contributing to faster computation.

The presented results established a new benchmark for NBC and represent a computational scale beyond what has been achieved by state-of-the-art DNA computing, which to date has solved a 20-variable 3-SAT problem ^20^. While the DNA computing technique exhaustively explored the full solution space of over one million truth assignments, it also required extensive processing steps, including sequential capture-and-release across 24 clause modules, followed by PCR amplification and gel-based readout to isolate and identify the correct solution. In contrast, NBC devices in this work solved Exact Cover instances with solution spaces on the order of 16 to 33 million possibilities, an order of magnitude larger than the DNA computing study. This scaling is achieved through the use of optimization strategies such as bit mapping optimization, entrance optimization, and reverse exploration (see Korten et al. 2021 for details of the optimization algorithm ^13^), which reduce network complexity while still guaranteeing correct solution identification. This difference in approach allows NBC to reach far larger problem spaces while maintaining practical device sizes and operational simplicity, by reducing the solution space that needs to be exhaustively explored. Although larger problems in NBC require a proportionally greater number of microtubules, this constraint can be mitigated by the ability to continuously supply microtubules during computation. Furthermore, producing large numbers of microtubules is far cheaper, because they self-assemble, allowing advances such as on-chip microtubule multiplication ^21^, which provide a means to sustain agent numbers during computation, further supporting scalability.

With the optimized junctions, for an overall success of 95%, networks sizes with at least 900 junctions can be solved in theory, however, in practice, this is currently limited by landing errors. These errors originated from microtubule landing within the computational network at unintended positions. In our experiments, despite the practically error-free junction operation, microtubules were still observed at incorrect exits while solving the 24-set and 25-set instances of the Exact Cover problem. 127 out of 2,667 filaments exhibited such behavior, which are attributed to landing errors. Although these errors did not significantly affect identification of the correct solution, they highlight the importance of controlling filament loading in future systems. To address this, we propose integrating NBC devices with microfluidic systems. Flow-focusing can confine microtubules to specific loading zones, steering them away from computation regions, making this a viable strategy for minimizing landing errors and improving system scalability ^17^. Moreover, achieving a level of parallelism that allows NBC to outperform conventional electronics in time-to-solution will require further progress in key areas such as programmability, scalability, and readout technology ^22–24^. Such techniques would remove the reliance on bulky, high-resolution imaging systems, and enable deployment of denser, complex, reusable, programmable NBC networks capable of solving multiple problem instances.

In summary, this study demonstrates a significant advancement in scalability of network-based biocomputation. By practically eliminating junction errors and validating the solution of large-scale problem instances representing over 30 million solutions we establish a new benchmark for molecular computing. Continued development in microfluidic integration, reconfigurable architectures, compact encoding strategies, and electrical readout platforms will be essential to realizing NBC’s potential as a viable, high-performance computing paradigm.

## Acknowledgements

This work was primarily supported and funded by the European Union’s Horizon 2020 research and innovation program under grant agreement no. 732482 (Bio4Comp). We further thank Corina Bräuer for technical support and all members of the Diez laboratory as well of the Bio4Comp consortium, in particular Hillel Kugler, Alf Mansson, Jingyuan Zhu, and Pradheebha Surendiran, for valuable scientific discussions. We also thank Richard Best for his help with the atomic force microscopy experiments. Additionally, we acknowledge support from the Dresden International Graduate School for Interdisciplinary Life Sciences (DIGS-ILS) and the Deutsche Forschungsgemeinschaft (DFG) within GRK 2767: Supracolloidal Structures: From Materials to Optical and Electronic Devices (Project No. 451785257).

## Author Contributions

D.R., H.L., T.K., and S.D. conceived the study. T.K. designed and C.R.M. nanofabricated the computation networks. E.C.V.R. and T.K. performed microtubule−kinesin experiments and analyzed the respective data. B.N., R.L., and C.R. contributed to testing and optimizing microtubule motility in the nano-channels. E.C.V.R., T.K., and S.D. wrote the manuscript with contributions from the other authors. All authors discussed the results and approved the final version of this manuscript.

## Methods

### Materials

All reagents were purchased from Sigma-Aldrich unless otherwise specified.

### Microtubule and Kinesin Preparation

Microtubules were polymerized from porcine brain tubulin (purified using established protocols ^25^), labeled with rhodamine in BRB80 buffer (80 mM PIPES, 1 mM EGTA, 1 mM MgCl_2_, pH 6.9), supplemented with 5 mM MgCl_2_, 1 mM MgGTP, and 5% DMSO. Polymerization was carried out at 37 °C for 1 hour using a 4 mg/mL tubulin mixture. The resulting microtubules were stabilized with 10 mM Taxol in BRB80 buffer and stored at room temperature. All experiments were conducted within 24 hours of polymerization. Full-length Drosophila melanogaster kinesin-1 motor proteins were expressed in insect cells and purified as described previously ^26^.

### Fabrication and assembly of Network-Based Biocomputing Devices

NBC networks were patterned on polymethyl methacrylate (PMMA)-coated 150 mm highly p-doped silicon wafers with a 75 nm silicon oxide layer. The substrates were cleaned using a standard RCA protocol to remove organic and metallic contaminants: a 1:1:5 mixture of NH_4_OH, H_2_O_2_, and deionized water (SC-1) followed by a 1:1:6 mixture of HCl, H_2_O_2_, and deionized water (SC-2). Both steps were conducted at 75 °C for 2 minutes. A 75 nm thermal SiO_2_ layer was then grown at 1000 °C in an oxygen and HCl-rich atmosphere, followed by annealing at 700 °C under nitrogen for 30 minutes to improve oxide quality. A post-annealing step at 200 °C for 30 minutes was performed to enhance PMMA adhesion.

Electron-beam resist (PMMA, Allresist AR-P 679.04) was spin-coated at 1700 rpm to achieve a thickness of ∼440 nm and hard-baked at 180 °C for 5 minutes. Devices fabrication proceeded in two steps (see Supplementary Figure S2): first, structures were patterned on PMMA using an VISTEC SB25, VISTEC Electron Beam GmbH, Jena, Germany operated at an accelerating voltage of 50 kV, an exposure dose of 650 µC/cm^2^. The electron beam (variable shaped beam) was raster-scanned according to the designed pattern using design in gds format generated by layout editor/K-Layout and transferred using ePLACE to machine compatible data. The patterned structures were then developed in a 1:3 mixture of MIBK and isopropanol for 60 seconds at room temperature, and rinsed with isopropanol and deionized water, and dried. Second, residual PMMA monolayers were removed from the channel floor by reactive ion etching (RIE) with CF_4_ plasma (20 sccm flow, 80 W RF power, 15-s exposure). Wafers were then cleaved into 10 mm × 10 mm chips for experimental use.

### Characterization of NBC devices

Scanning electron microscopy was conducted using a GAIA3 (TESCAN, Czech Republic) operated at an accelerating voltage of 5.0 kV and a working distance in the range of 9.93 to 10.02 mm. An InBeam SE detector was employed to obtain high-resolution images. Samples were directly imaged without sputter-coating any conductive layer.

The channel width was measured using atomic force microscopy (AFM) in non-contact mode. AFM measurements were performed in non-contact mode using a TI950 Nanoindenter platform (Hysitron/Bruker, USA). Scans were acquired over an area of 3 × 3 µm^2^ with a resolution of 512 × 512 pixels using a cantilever with a nominal tip radius of 7 nm. Topographic data were processed and analyzed using Gwyddion software.

### Experimental Validation of NBC devices

Flow cells were assembled by placing two parallel strips of parafilm (∼ 2 mm apart) on either side of the patterned region on the fabricated chips, followed by placement of a 22 × 22 mm^2^ Diphenyldimethoxysilane (DDS)-coated coverslip on top. The assembly was sealed by gentle heating for ∼20 seconds, producing a leak-proof channel. The assembled flow cells were perfused with Pluronic F127 in BRB80 buffer and allowed to incubate for 2 hours at room temperature to selectively passivate the PMMA surface, while leaving the SiO_2_ channel floor exposed for kinesin binding ^**14**^. Following this, the flow cell was perfused with 0.5 mg/mL casein in BRB80 buffer and incubated for 5 minutes. Subsequently, 5 μL of 4 nM kinesin-1 solution was introduced and incubated for 15 minutes to allow motor protein attachment. Finally, a motility buffer containing 1 mM Mg-ATP, 20 mM D-glucose, 20 μg/mL glucose oxidase, 10 μg/mL catalase, 10 mM DTT, and 10 μM Taxol in BRB80 was introduced, followed by rhodamine-labeled, taxol-stabilized microtubules.

### Image Acquisition

Gliding motility assays were imaged using a Nikon Eclipse Ni upright fluorescence microscope equipped with a CFI S Plan Fluor ELWD 20X long-distance, wide-field air objective and an ORCA-Fusion 14440 camera. Images were acquired and processed with NIS-Elements software (Nikon). CoolLED pE4000 was used as the light source, and a multiband filter (DAPI/FITC/Cy3/Cy5/Cy7 Penta-LED HC Filter, Semrock) was used to visualize the rhodamine-labeled microtubules. Exposure time was set to 500 milliseconds, and time-lapse images were recorded at 2-second intervals.

### Data Analysis and Statistical Methods

To evaluate the reliability of the solution in NBC, both landing errors and junction errors were quantitatively assessed. Landing errors were estimated using the equation:

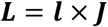

where*l* is the landing rate per junction, and *J* is the number of junctions

The junction error rate, E was defined as:

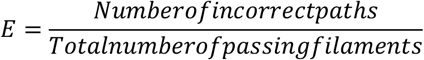

The probability that a microtubule agent follows the correct path across *X* junctions scales as (1 – *E*) ^*x*^. Junction error rates are reported as percentages in the Results and Discussion to represent junction performance.

To evaluate whether the filament distribution at the network exits reliably indicated a correct solution (e.g., an exact cover), a four-step statistical framework based on our previous work was employed ^14^. This approach integrates empirical error estimates with statistical modeling to determine whether observed filament counts could be attributed to accurate computation rather than random fluctuations. Total filaments counts affected by guidance errors were estimated by combining experimentally measured junction error rate with landing error estimates from Monte Carlo simulations. These data were normalized relative to the number of network exits, enabling robust confidence assessments to distinguish between correct and wrong exits.

The maximum projection images of frequented microtubules paths were made using Fiji ImageJ software, and the plots were plotted using Origin Pro 2024b. Linear regression and statistical analysis were performed in MATLAB.

## Supplementary Figures

**Figure S1.**
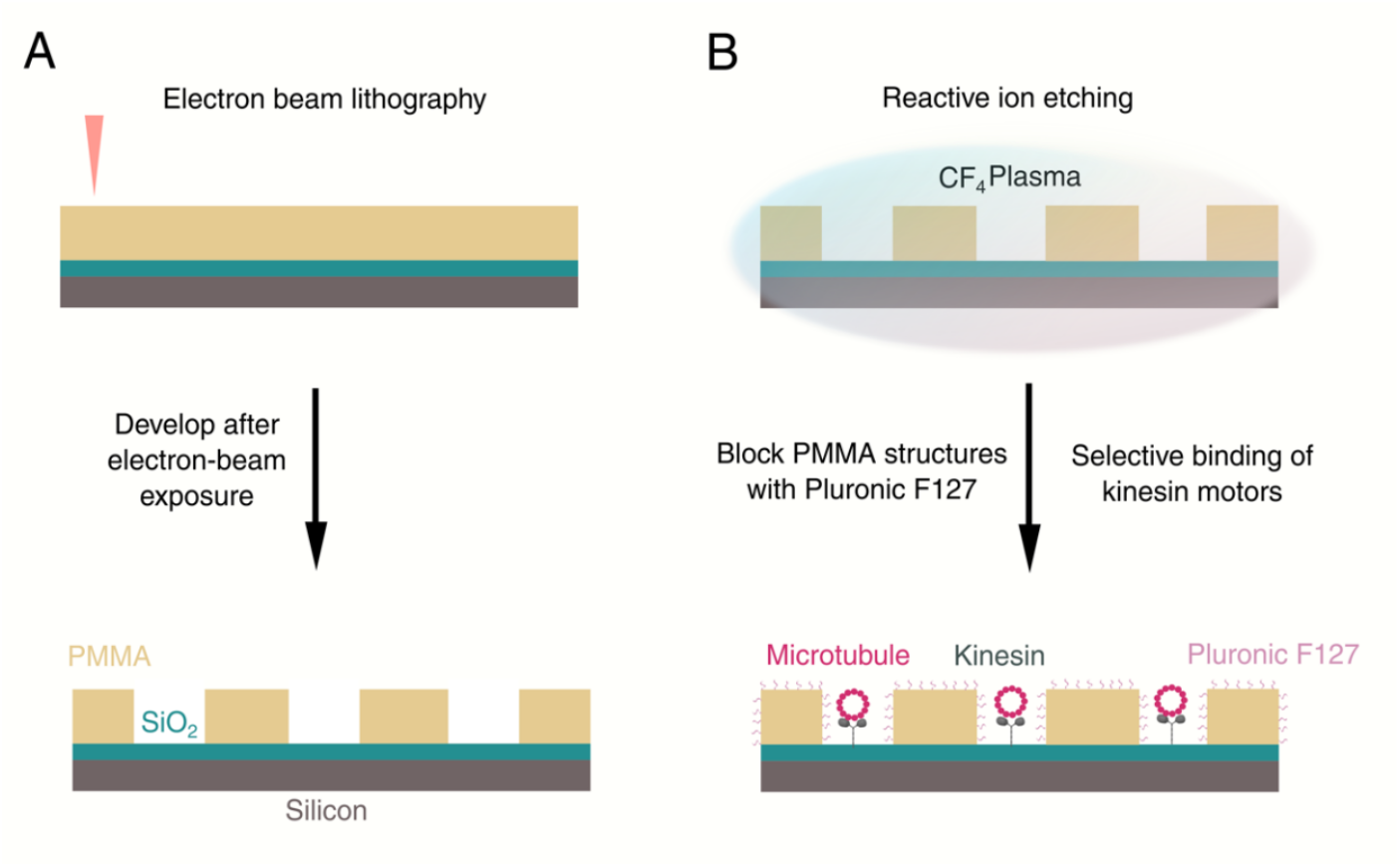
Schematic of modified fabrication protocol for patterning sub-200 nm PMMA structures. (**A**) PMMA coated Silicon/SiO_2_ substrate was patterned with electron beam lithography. PMMA layer is shown in yellow, SiO2 layer is shown in green and silicon substrate is shown in brown. Following electron beam exposure, the structures are developed with wet etching. (**B**) The developed structures are subjected to reactive ion etching in the presence of CF_4_ plasma. This is followed by blocking the PMMA walls with Pluronic F127. Blocking the PMMA surface with Pluronic F127 prevents kinesin from binding to the PMMA surfaces and selectively bind to the SiO_2_ surface on the channel floor.

**Figure S2.**
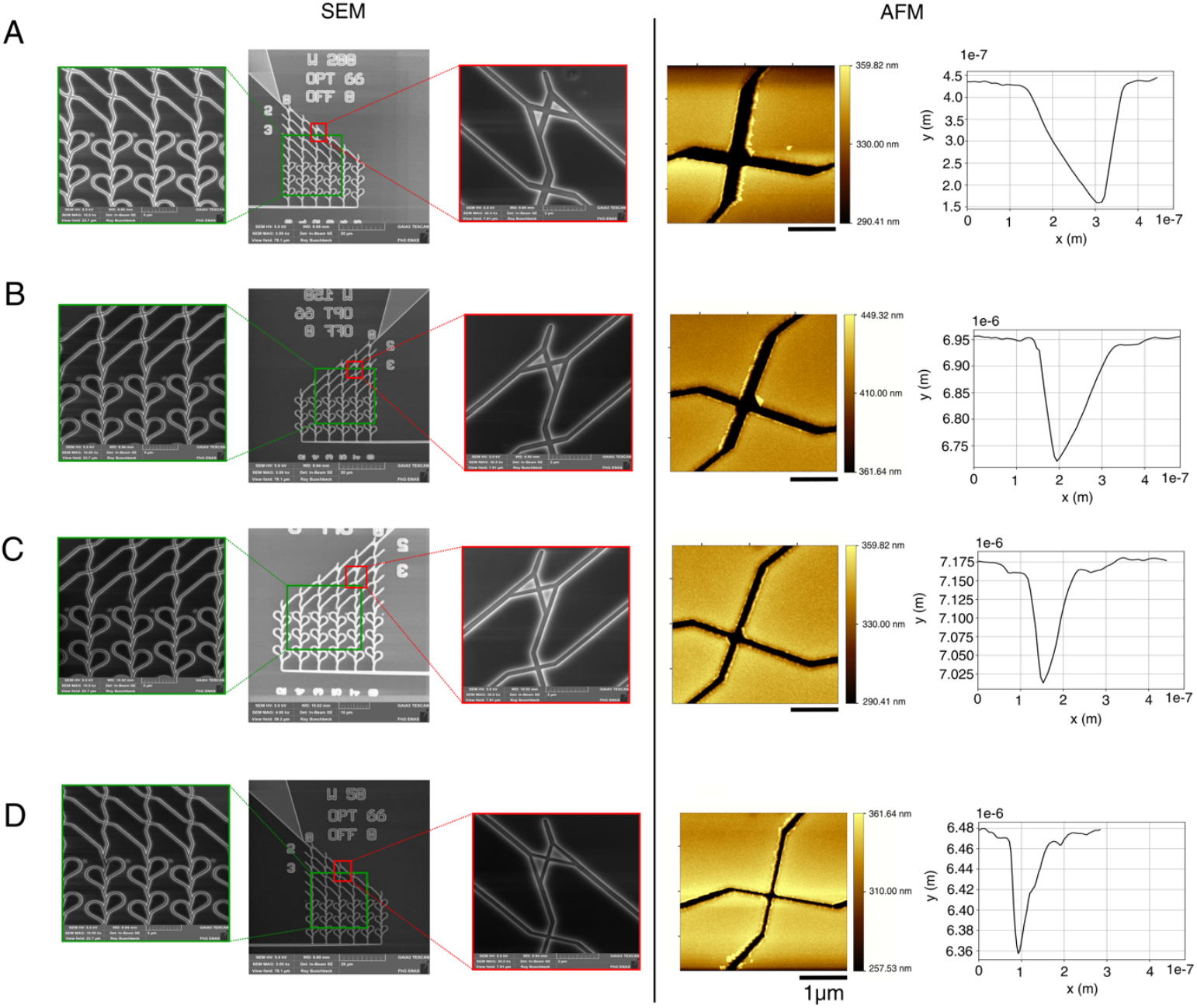
Physical characterization of junction geometries with varying widths. Scanning electron microscopy (SEM) and atomic force microscopy (AFM) analysis of pass junction geometries with varying channel widths. (**A**) 200 nm, (**B**) 150 nm, (**C**) 100 nm, and (**D**) 50 nm. Left panels (SEM): Low- and high-magnification images show the overall layout of the fabricated networks (green insets) and individual junctions (red insets). Right panels (AFM): Topographic maps (Scale bar = 1 µm) and corresponding cross-sectional profiles confirm the fabricated channel dimensions. Based on AFM measurements, the measured widths for the designed junctions were 231 nm ± 16 nm (n = 5) (mean ± standard deviation, n = 5), 170 nm ± 6 nm (n = 5), 117 nm ± 9 nm (n = 5), and 71 nm ± 8 nm, respectively.

**Figure S3.**
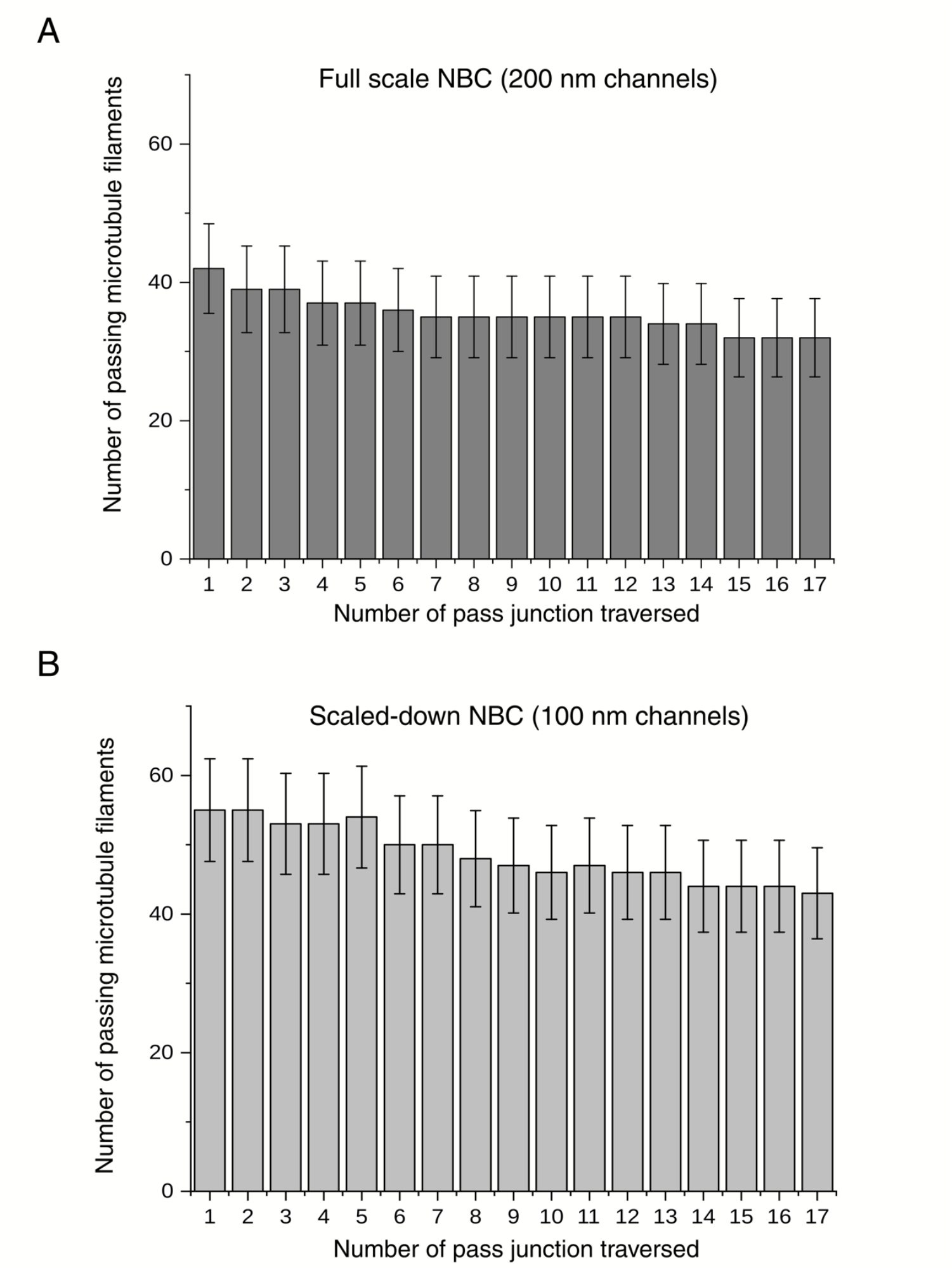
Detachment rate of microtubules per junction. (**A**) Bar plots showing the number of microtubules crossing each junction and the number of junctions traversed on the full-scale NBC with 200 nm channels. 10 out of 42 microtubules filaments detached before crossing 17 pass junctions. Error bars denote counting errors. (**B**) Bar plots showing the number of microtubules crossing each junction and the number of junctions traversed on the scaled-down NBC with 100 nm channels. 11 out of 55 microtubule filaments detached before crossing 17 pass junctions. Error bars denote counting errors.

